# CLIPreg: Constructing translational regulatory networks from CLIP-, Ribo- and RNA-seq

**DOI:** 10.1101/2021.12.06.470871

**Authors:** Baptiste Kerouanton, Sebastian Schäfer, Lena Ho, Sonia Chothani, Owen J. L. Rackham

## Abstract

**Motivation:** The creation and analysis of gene regulatory networks have been the focus of bioinformatic research and underpins much of what is known about gene regulation. However, as a result of a bias in the availability of data-types that are collected, the vast majority of gene regulatory network resources and tools have focused on either transcriptional regulation or protein-protein interactions. This has left other areas of regulation, for instance translational regulation, vastly underrepresented despite them having been shown to play a critical role in both health and disease.

**Results:** In order to address this we have developed CLIPreg, a package that integrates RNA, Ribo and CLIP-sequencing data in order to construct translational regulatory networks coordinated by RNA-binding proteins. This is the first tool of its type to be created, allowing for detailed investigation into a previously unseen layer of regulation.

**Availability and implementation:** CLIPreg is available at https://github.com/SGDDNB/CLIPreg.

## 1 Introduction

Cells are governed by a complex network of interactions that coordinate each layer of gene expression and ultimately determine the function of the cell. Because of the interconnected nature of this network it is important that we do not study each gene in isolation but also consider their interactions with and effects on all of the other genes. As a result, much attention has been placed on the experimental and computational methods that can be used to observe, understand and intervene with these networks as a means to better understand biology (see (Ideker and Nussinov, 2017) for an overview) and to find new treatments to disease (Barabási *et al*., 2010). To date, the vast majority of these methods have focused on either the transcriptional regulation (i.e. Transcription factor - DNA interactions) or protein-protein interactions. The reason for this is that the tools available for assaying these have become more widespread and easier to access and not as a result of their relative importance to the overall gene regulation of a cell. For instance, in a recent paper it was shown that over one-third of all changes that occur in fibrosis have an element of translational regulation (Chothani, Schäfer, *et al*., 2019) and dysregulation of translation has been shown to impair erythropoiesis (Alvarez-Dominguez *et al*., 2017) and lead to cancer metastasis (Micalizzi *et al*., 2021). To address this we have developed the CLIPreg package that combines data on translational regulation (via RNA sequencing [RNA-seq] and Ribosome profiling [Ribo-seq]) with data on protein-RNA interactions (via Cross-linking immunoprecipitation and sequencing [CLIP-seq]) to construct a translational regulation network.

## 2 CLIPreg Workflow and Outputs

Here, we describe the key features and usage of the CLIPreg R package as well as the available visualisations to help understand translational regulation by RNA-binding proteins (RBPs) occurring in a system of interest. To demonstrate this, we provide an example analysis of RBP driven translational regulation in fibroblast activation (based on a recent publication (Chothani, Schäfer, *et al*., 2019)). However, this package can be applied to any given matched Ribo- and RNA-seq from two or more conditions (e.g. WT vs control or a time-series). CLIP-seq data can either be imported from an existing database (POSTAR (Hu *et al*., 2017) and ENCODE (Van Nostrand *et al*., 2020)) or provided directly by the user.

### 2.1 Key features of CLIPreg

The purpose of CLIPreg is to automate the identification of key RBPs regulating translation through the integration of RNA- Ribo- and CLIP-seq data. The package has three stages (1) Data preparation, (2) Data integration and (3) Data visualisation and analysis (outlined in Figure 1A). The combination of these steps allows a user to generate hypotheses concerning the role of RBPs in their system of interest.

**Fig. 1.**
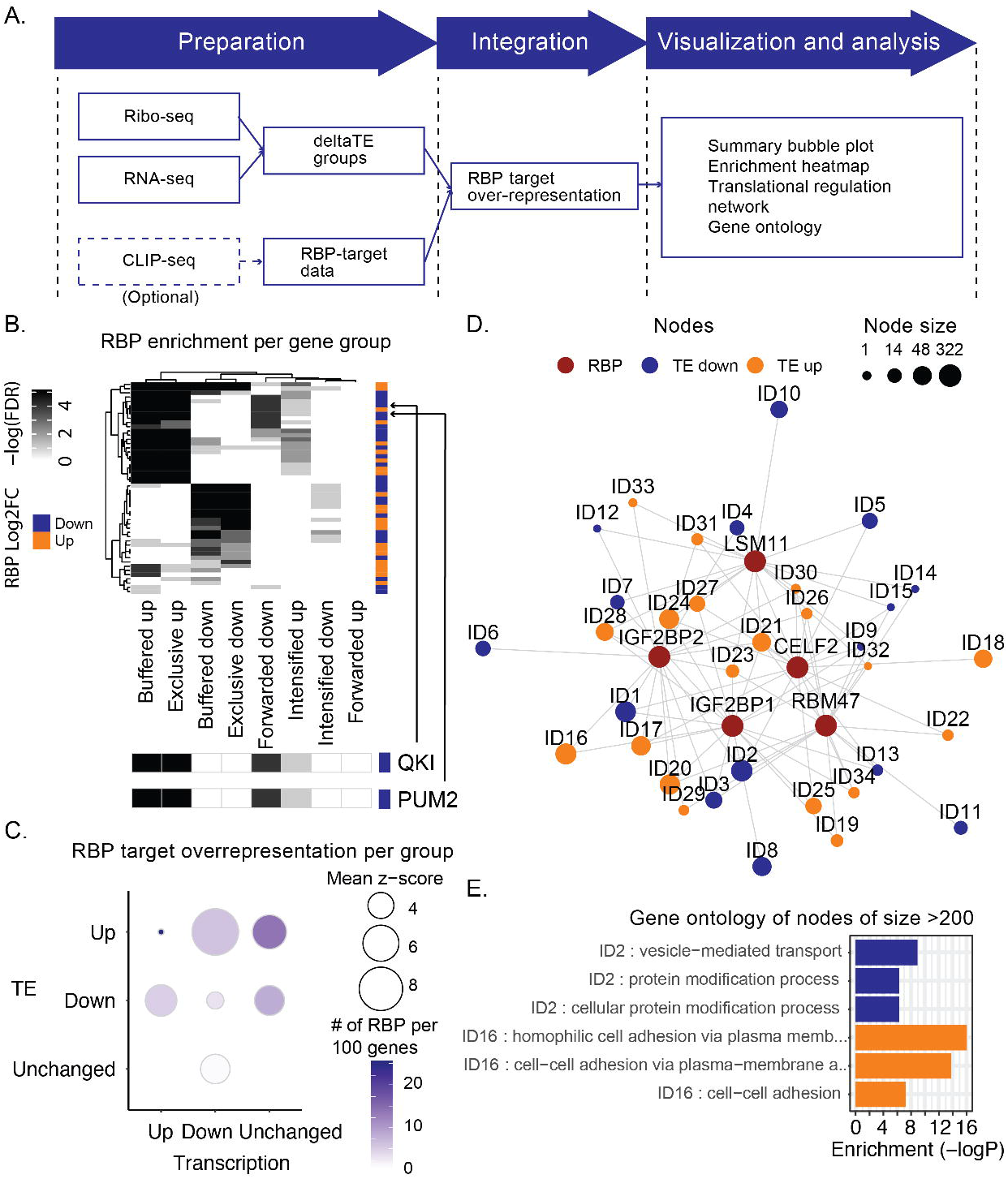
(a) CLIPreg workflow: Step 1, Data preparation: Ribo-seq and RNA-seq data are used to determine translational efficiency changes using deltaTE (Chothani, Adami, et al., 2019) and CLIP-seq data is used to determine RBP-target relations. Step 2, Data integration: RNA-binding protein (RBP)-target over-representation test to identify key RBPs regulating translation. Step 3, Data visualisation: Visualisation and analysis. (b) Heatmap of RBP enrichment per gene group. -log(P) denotes the false discovery rate. RBP Log2FC denotes RBP’s Ribo-seq log_2_ fold change. Orange: RBP upregulated; Blue: RBP downregulated. Quaking (QKI) and Pumillo-2 (PUM2) highlighted for detailed view. (c) Bubble plot showing RBP enrichment per regulation group. (d) RBP Translational regulation network. Red: RBP; Blue: Targets with upregulated translational-efficiency (TE); Orange: Targets with downregulated TE. (e) Gene ontology of RBP regulated gene sets as labelled in the network. ID2 and ID16 groups shown as examples.

In the dataset preparation step, two separate sources of information must be supplied: (a) Groups of genes that are known to be translationally regulated via the application of the DeltaTE package (Chothani, Adami, *et al*., 2019). In total there are eight of these groups (which we refer to as DeltaTE groups), each of which capture sets of genes with similar translational regulation profiles, for instance, the buffered down group (referring to genes that increase transcriptionally but have translational buffering) or intensified up group (referring to genes that are up-regulated both transcriptionally and translationally) (b) Known interactions between RBPs and mRNAs. The package by default uses interaction data derived from the POSTAR (Hu *et al*., 2017) and ENCODE (Van Nostrand *et al*., 2016) databases but a user can also use their own data. The result of this step is that all required data is formatted and ready for analysis.

In the dataset integration step, the CLIPreg R package looks for the over-representation of RBPs in each of the DeltaTE groups. This is done by calculating the empirical p-value of the frequency of interactions between any given RBP and each individual DeltaTE group by comparing the number of observed interactions with a null distribution generated from repeated shuffling (n = 100,000 iterations) of the RBP-mRNA interactions. This results in a set of RBP regulators enriched in each group, together with an associated confidence corrected for multiple testing (false discovery rate) and effect size (z-score) for each RBP.

In the data visualisation and analysis steps, a number of summary plots and analyses are produced. (a) A heatmap is generated that links the set of RBPs to each DeltaTE group (see Figure 1B). The purpose of this plot is to identify patterns in RBPs that are enriched in similar modes of translation regulation, for instance buffering down and exclusive down (referring to genes where the mRNA level doesn’t change but the rate of translation decreases) are both similar modes of translational regulation so it should be expected that similar RBPs coordinate this regulation. In order to make this plot as information rich as possible, RBPs are clustered so that RBPs with similar translational roles are close together. (b) To summarise the role of RBP-regulation in each of the DeltaTE groups, a bubble plot is generated to show the average enrichment and percentage of RBPs in each of the eight groups. Typically we don’t expect to see RBPs playing a role in transcriptional regulation (i.e. those groups where translational efficiency (TE) is unchanged) and this can be easily investigated in this plot. (c) A translational regulatory network is constructed and visualised showing how RBPs are predicted to regulate different groups of genes. For simplicity, genes are grouped into regulated gene sets (RGSs) according to which regulators they are connected to and uniquely labelled (eg ID1). This allows a user to visualise which RBPs appear to regulate the same genes. (d) For each RGS, a gene ontology analysis using topGO R package is performed based on its members to find any overrepresented gene functions. These can be visualised in order to guide a user towards a putative function of the RBP regulation.

### 2.2 Example Usage of CLIPreg

To demonstrate ClipReg we applied it to data from a recent publication where matched Ribo and RNA sequencing was generated from cardiac fibroblasts treated with Transforming growth factor beta 1 (TGFB1) as a model of fibrosis (Chothani, Schäfer, *et al*., 2019). The DeltaTE package was run, resulting in the identification of 4680 genes that were either transcriptionally and/or translationally regulated, each of which was placed into one of the eight DeltaTE groups. For each DeltaTE group, CLIPreg was used to calculate the enriched RBPs based on targets, identifying 50 RBPs enriched in at least one DeltaTE group. The significance of these enrichments is summarized for each RBP across the eight DeltaTE groups in Figure 1B, showing that there appears to be two main clusters of RBP regulation, one that is enriched in buffered and exclusive down genes (i.e. those where the rate of protein abundance is repressed translationally) and one that is enriched in exclusive and buffered up genes (i.e. those where the rate of protein abundance is enriched translationally). The DeltaTE groups where RBPs seem to be most active are identified in Figure 1C, showing that the buffered up group appears to have the strongest translation regulatory effect from RBPs, with the mean z-score of the observed frequency compared to the null distribution being ~8. Taken together, the heatmap and bubble plots reveal the importance of the RBPs across deltaTE groups. A detailed network analysis of the top 5 RBPs with largest fold changes are shown in Figure 1D, showing that there are 29 RGSs amongst these 5 RBPs. A GO analysis of these RGSs shows that one (ID2) is enriched for vesicle mediated transport and protein modification process whilst another (ID16) is enriched for genes involved in cell-cell adhesion.

### 2.3 Basic usage

The only required data from the user to run this package are the Ribo and RNA-seq count files that can be used in DeltaTE or the output of DeltaTE directly. The package contains one function for the enrichment analysis (run_CLIPreg) and functions for each plot. A detailed tutorial can be found in the Github repository but in general the package contains functions for each of the data preparation, dataset integration, data visualisation and analysis steps as follows

~~~
#STEP 1: data preparation step
CLIPreg_example=Load_example()
#STEP 2: data integration step
results=run_CLIPreg(input_data=CLIPreg_example, is.example=T)
#STEP 3: data visualisation and analysis step
dir.create(“Results_CLIPreg”)
Visualise(results=results, folder=“Results_CLIPreg”)
~~~

The output of the package is a directory containing 5 files: A .RData file that contains the output from the run_CLIPreg function and four pdf with the figures illustrated in Figure 1.

## 3 Discussion

The role of translation regulation has remained understudied until now due to the relative difficulty in obtaining data that can help uncover this layer of regulation. As the availability of matched Ribo- and RNA-seq data together with large repositories of CLIP-seq data increases, the need for methods to analyse these to generate testable hypotheses also grows. Existing attempts to do this have relied on protein-protein interaction information [eg (Wu *et al*., 2010)] or have focused on post-translational modifications to proteins [eg (Huang *et al*., 2018)] rather than the direct regulation of translation. As such, this tool will allow constructing translational regulatory networks, opening up the identification and therapeutic targeting of RBPs as a means to block or reverse disease onset or to better understand the role of translation in the establishment and maintenance of cell fate.

## Author Contributions and Funding

O.J.L.R, B.K and S.C conceived and designed the package with input from S.S and L.H. B.K. wrote the package with input from S.C. O.J.L.R managed the project with input from L.H and S.S and S.C. This work was supported by a Singapore National Research Foundation grant [NRF-CRP20-2017-0002]. Conflict of Interest: none declared.

## Notes

https://github.com/SGDDNB/CLIPreg

### Competing Interest Statement

The authors have declared no competing interest.

