## Supplementary file 1 for "CLIPreg: Constructing translational regulatory networks from CLIP-, Ribo- and RNA-seq"

### CLIPreg

The goal of CLIPreg is to discover key RBP regulators in different datasets. It combines CLIP-seq with RNA- and RIBO-seq to calculate enrichment of RBP and generate plots for publications.

### Installation

#### Check and install required packages

Users may use following codes to check and install all the required packages.

```
list.of.packages <- c("ggplot2","grid","doParallel","foreach","data.table","fastma

## for package "ggplot2", "pheatmap", "grid", "doParallel", "foreach", "data.table
new.packages <- list.of.packages[!(list.of.packages %in% installed.packages()[,"Pa
if(length(new.packages)) install.packages(new.packages)

## for package "topGO", "ALL", "ggnet", "ComplexHeatmap","org.Hs.eg.db"
if (!requireNamespace("BiocManager", quietly = TRUE)) install.packages("BiocManage
new.packages <- list.of.packages[!(list.of.packages %in% installed.packages()[,"Pa
if(length(new.packages))  BiocManager::install(new.packages)
if("ggnet"%in%new.packages)  devtools::install_github("briatte/ggnet")
install_version("network", version = "1.16.1", repos = "http://cran.us.r-project.o
if("ComplexHeatmap"%in%new.packages)  devtools::install_github("jokergoo/ComplexHe
```

#### Install CLIPreg

The source code of CLIPreg can be installed from GitHub with:

```
devtools::install_github("SGDDNB/CLIPreg")
```

CLIPreg requires 4 different inputs. A gene_groups input which is a dataframe containing geneID and the gene_groups given by DeltaTE output (ideally). The DeltaTE method can be found in the paper here: https://doi.org/10.1002/cpmb.108. It categorizes transcriptionally and/or translationally regulated genes into 4 categories: forwarded, intensified, exclusive and buffered, each category with an up or down direction. Summarized CLIP-seq data, POSTAR and ENCODE are pre-loaded in the package. RIBO lfc and TPM.

```
## load libraries
library(CLIPreg)
library(ggplot2)
library(ComplexHeatmap)
library(grid)
library(doParallel)
library(ggnet)
library(topGO)
library(GGally)
library(data.table)
library(stringr)
```

### Basic usage

For testing the package with the example data

```
Example=Load_example()
```

For using your own data, you must specify a folder that contains at least 3 txt files that are named: gene_groups.txt, ribo_lfc.txt and ribo_tpm.txt. For the format of those 3 files, please refer to Advance usage step 1.

```
Input_data=Load_input_files(folder = "path/to/folder")
```

Run the analysis with all default parameters.

```
results=run_CLIPreg(Example, is.example=T) # or run_CLIPreg(Input_data, is.example
```

Generate the visual output from the results of the analysis. The Visualise function will create 2 new files in the folder given: an Rdata file containing the list object from the run_CLIPreg function and a pdf file with the main figures.

```
dir.create("Results_CLIPreg")
Visualise(results=results,folder="Results_CLIPreg")
```

### Advance usage

#### Step 1: Input datasets

Let’s have a look at the gene_groups file from the example data. It consists of 2 columns containing the geneID and the gene groups for all the DE genes. This is optional, if you want to provide your own gene groups, go to next step.

```
data("gene_groups")
head(gene_groups)
#>              geneID   Gene_group
#> 1:        LOC728392 forwarded_up
#> 2:      WRB-SH3BGR forwarded_up
#> 3: ENSG00000162408 forwarded_up
#> 4: ENSG00000204859 forwarded_up
#> 5: ENSG00000142583 forwarded_up
#> 6: ENSG00000116649 forwarded_up
```

You can input your own gene groups by using the function load_gene_groups() and giving the file location of your file as input.

```
#gene_groups=load_gene_groups(gene_groups_file = "path/to/gene_groups_file.txt")
```

Load POSTAR and ENCODE RBP data. Those are 2 public datasets which are processed in order to have lists of vector. Each vector is named after 1 RBP and contains the geneID of all the targets of that RBP. Combine both data in a target list.

```
data("RBP_ENCODE")
data("RBP_POSTAR")
Targets=combine_targets(RBP_list1=RBP_ENCODE,RBP_list2=RBP_POSTAR,background=gene_
```

Load the fold change and identify the RBPs. If you have your own then provide your own data.

```
# load fold change and tpm. optional if you want to use the input data
data("ribo_lfc")
data("tpm_ribo")

head(ribo_lfc)
#>                           geneID IDENTIFIER FoldChange
```

```
#> ENSG00000000003 ENSG00000000003     TSPAN6  0.3669041
#> ENSG00000000419 ENSG00000000419       DPM1  0.1026692
#> ENSG00000000457 ENSG00000000457      SCYL3 -0.4238081
#> ENSG00000000971 ENSG00000000971        CFH -0.2162611
#> ENSG00000001084 ENSG00000001084       GCLC -0.1003380
#> ENSG00000001461 ENSG00000001461     NIPAL3 -0.3866424
#head(tpm_ribo)

# If you want to input your own data
#load_ribo_lfc(ribo_lfc_file = "ribo_lfc_file")
#load_ribo_tpm(ribo_tpm_file = "ribo_tpm_file")
```

#### Step 2: Data integration and analysis

Run the enrichment analysis using CLIPreg() function. This takes several minutes. If you want to have a look at the example results skip this step.

```
# The CLIPreg function requires a few minutes to run, save the data after running
# res_Postar=CLIPreg(RBP_data=RBP_POSTAR,gene_groups=gene_groups)
# res_Encode=CLIPreg(RBP_data=RBP_ENCODE,gene_groups=gene_groups)
# save(res_Encode,file="Res_RBP_Encode.RData")
# save(res_Postar,file="Res_RBP_Postar.RData")
```

If you want to get the results directly you can load it by using the example data results. The output of CLIPreg() is a list of dataframes. One dataframe per gene group containing the RBP and statistical information calculated during the analysis such as p-value and z-score.

```
data("res_Encode")
data("res_Postar")
head(res_Encode[[1]])
#>       RBP real_overlap simulated_overlap_mean simulated_overlap_sd          z
#> 1 ZC3H11A          268              282.41101            12.590167 -1.1446242
#> 2    GNL3           14               15.73466             3.306621 -0.5246020
#> 3  HNRNPM          345              396.49065            14.077142 -3.6577488
#> 4   RBM15          461              476.04563            14.839723 -1.0138754
#> 5   DDX24          370              406.79695            14.246241 -2.5829235
#> 6   XRCC6          142              146.71529             9.573704 -0.4925252
#>      pval padj
#> 1 0.86594    1
#> 2 0.63920    1
#> 3 0.99986    1
#> 4 0.83574    1
```

```
#> 5 0.99474    1
#> 6 0.66889    1
```

Then we want to combine POSTAR and ENCODE to work with only one dataframe and only keep RBPs that are significant in at least one gene group.

```
res=CLIPreg::combine(res1=res_Encode,res2=res_Postar,FDR=0.1)
```

Extract the RBP LFC from the RIBO_LFC and keep only detected RBPs in res

```
# Change of RBPs
rbp_lfc=rbp_change(res=res,ribo_lfc=ribo_lfc)

# Cure res data by removing RBPs that are not in the rbp_lfc dataframe
res=cure_res(res=res,rbp_lfc=rbp_lfc)
```

#### Step 3: Visualisation

Generate and save heatmap to pdf. The heatmap represents the -logP of each RBP for each gene group. The blue RBPs are downregulated and the orange RBPs are upregulated. Only RBPs that are significant in at least one gene group are shown.

```
# Heatmap of RBP scores

HeatmapRBP(res=res,rbp_lfc=rbp_lfc)
```

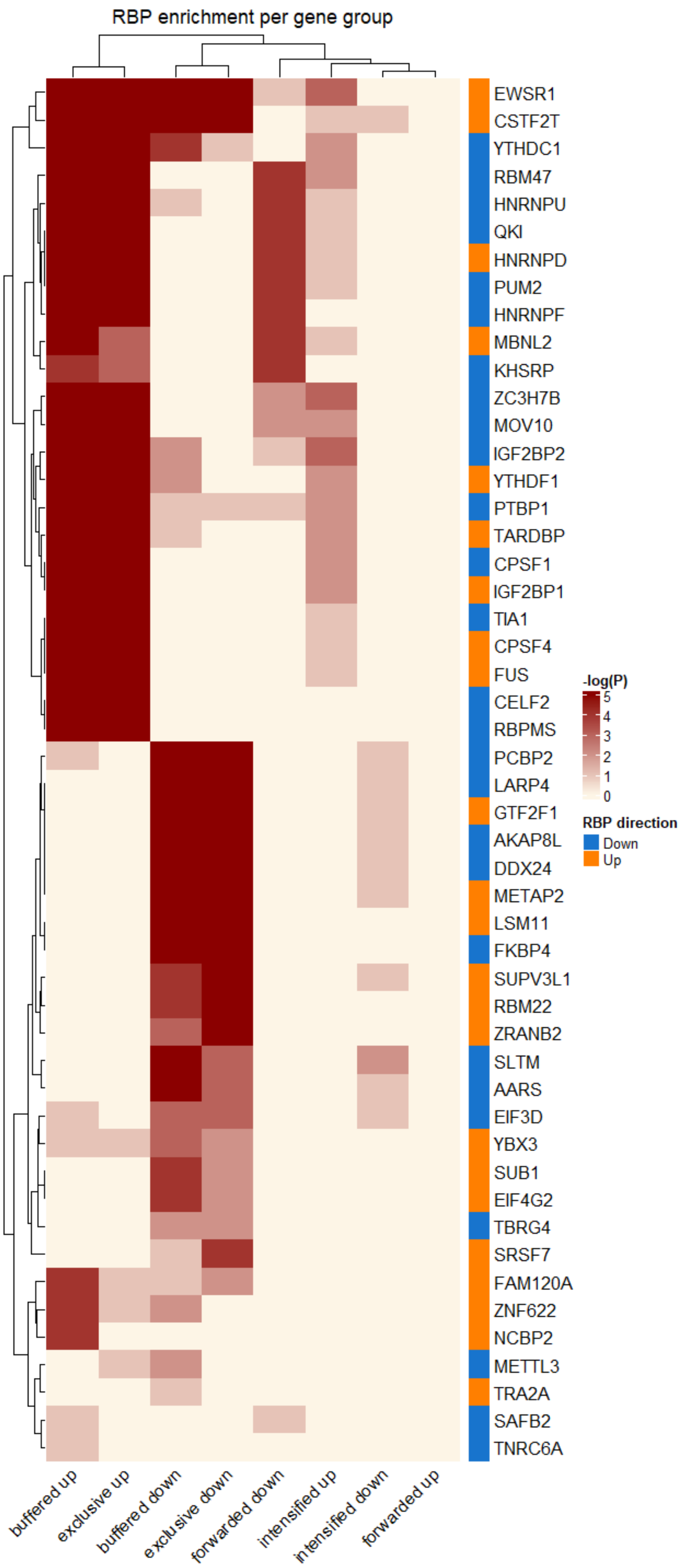

RBP enrichment per gene group
EWSR1
CSTF2T
YTHDC1
RBM47
HNRNPU
QKI
HNRNPD
PUM2
HNRNPF
MBNL2
KHSRP
ZC3H7B
MOV10
IGF2BP2
YTHDF1
PTBP1
TARDBP
CPSF1
IGF2BP1
TIA1
CPSF4
FUS
CELF2
RBPMS
PCBP2
LARP4
GTF2F1
AKAP8L
DDX24
METAP2
LSM11
FKBP4
SUPV3L1
RBM22
ZRANB2
SLTM
AARS
EIF3D
YBX3
SUB1
EIF4G2
TBRG4
SRSF7
FAM120A
ZNF622
NCBP2
METTL3
TRA2A
SAFB2
TNRC6A
-log(P)
5
4
3
2
1
0
RBP direction
Down
Up
buffered up
exclusive up
buffered down
exclusive down
forwarded down
intensified up
intensified down
forwarded up

```
# Plot the heatmap with updated colors for the RBPs

# Save the heatmap
# e=HeatmapRBP(res=res,rbp_lfc=rbp_lfc)
# location="Heatmap_fibroblasts.pdf"
# n=length(e$tree_row$order)
# pdf(location,8,3+n*0.15)
#
# dev.off()
```

A bubble plot can be generated only if the gene groups are the output of DeltaTE program.

```
# Bubble plot gene_groups if gene_groups are from DeltaTE. FDR has to be lower or
BubbleRBPs(res = res,gene_groups = gene_groups,rbp_lfc = rbp_lfc,FDR=0.1)
```

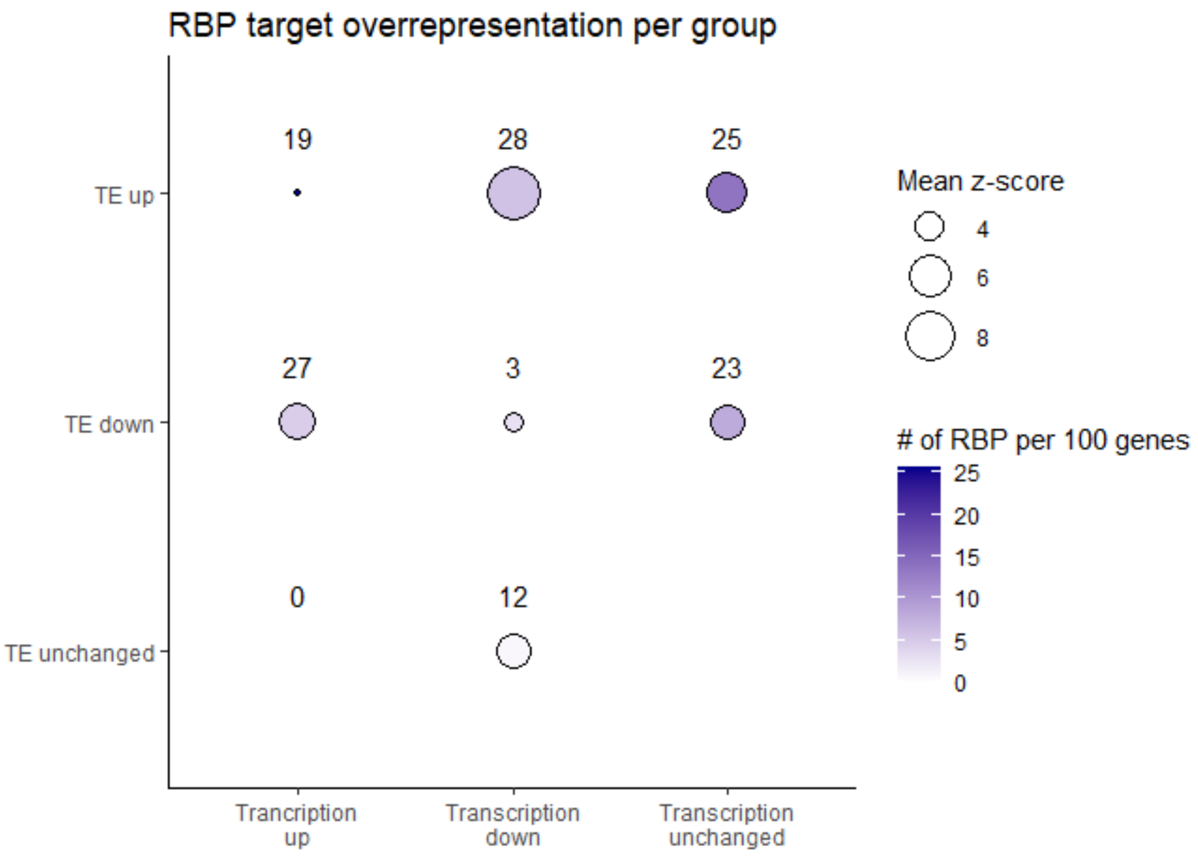


From the results, the user can choose a number of RBP to draw the network for by n. This will pick the n most changing RBPs.

```
# Draw network
```

```
Draw_network_by_group(rbp_lfc=rbp_lfc,res=res,Targets=Targets,gene_groups=gene_gro
```

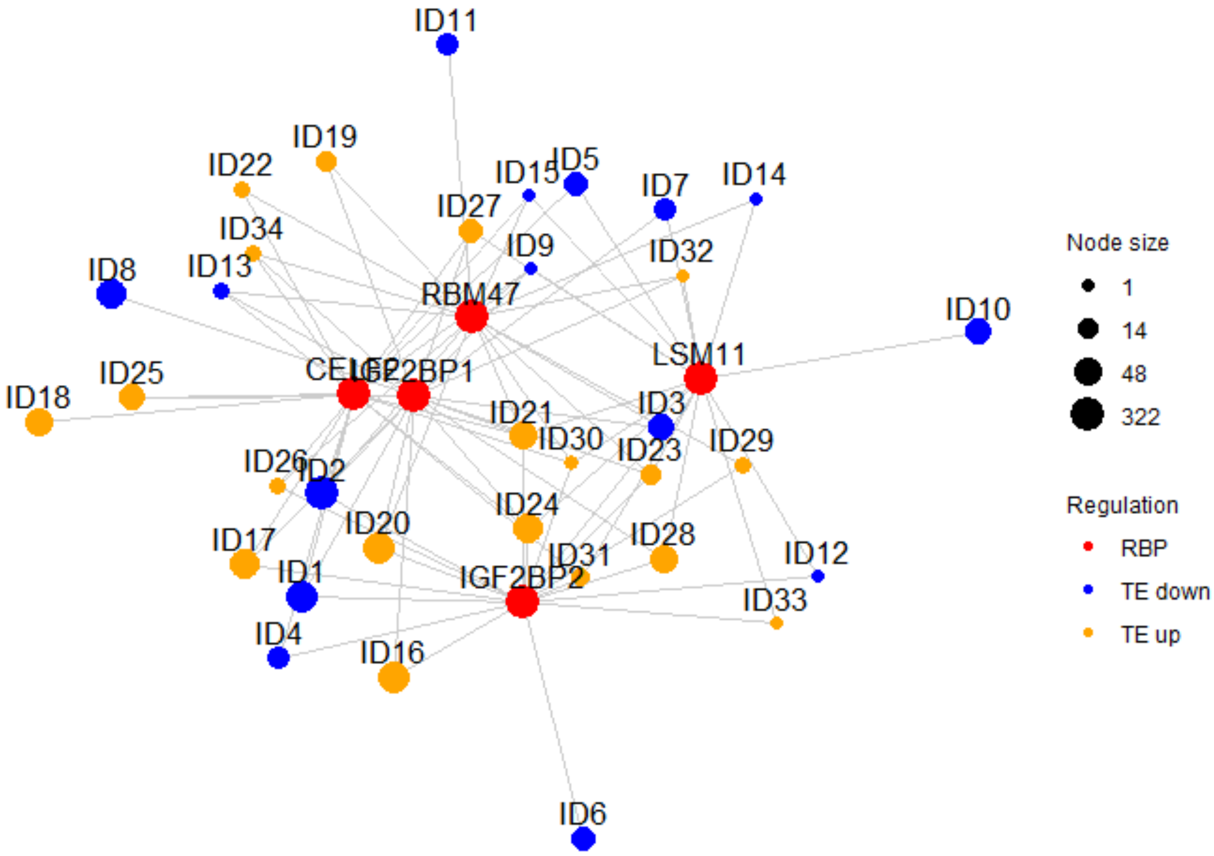


```
# plot GO
Plot_GO(rbp_lfc=rbp_lfc,res=res,Targets=Targets,gene_groups=gene_groups,n=5,
  tpm_ribo = tpm_ribo,th=200,GO_to_show=3,forwarded = F)
```

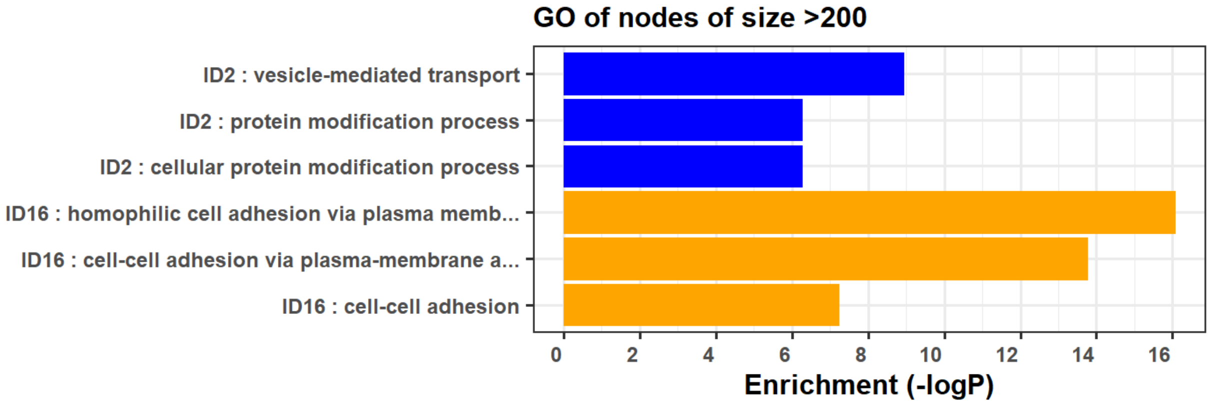


```
Plot_GO_node_name(rbp_lfc=rbp_lfc,res=res,Targets=Targets,gene_groups=gene_groups,
                  tpm_ribo = tpm_ribo,Nodes_to_keep=c(19,15),GO_to_show=3,forwarde
```

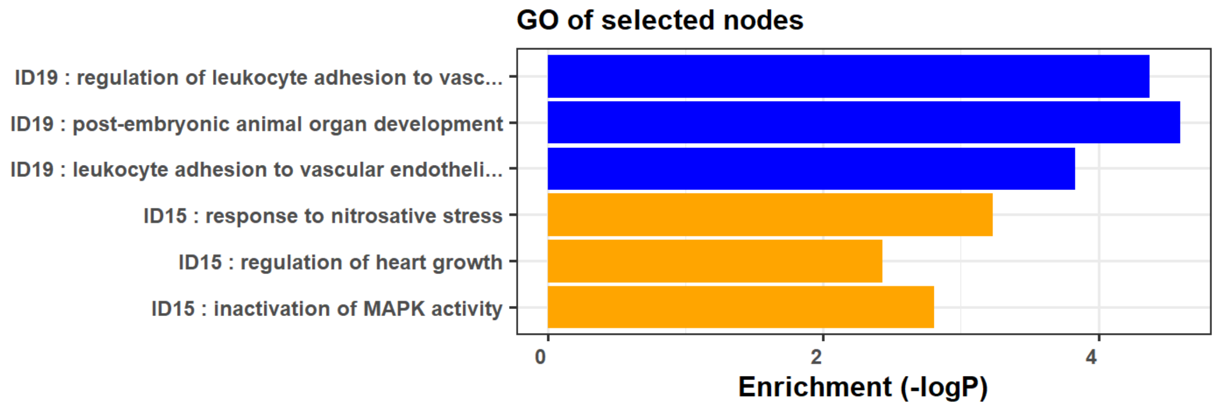


```
Plot_GO_RBP(rbp_of_interest="QKI",tpm_ribo = tpm_ribo,Targets=Targets,gene_groups=
```

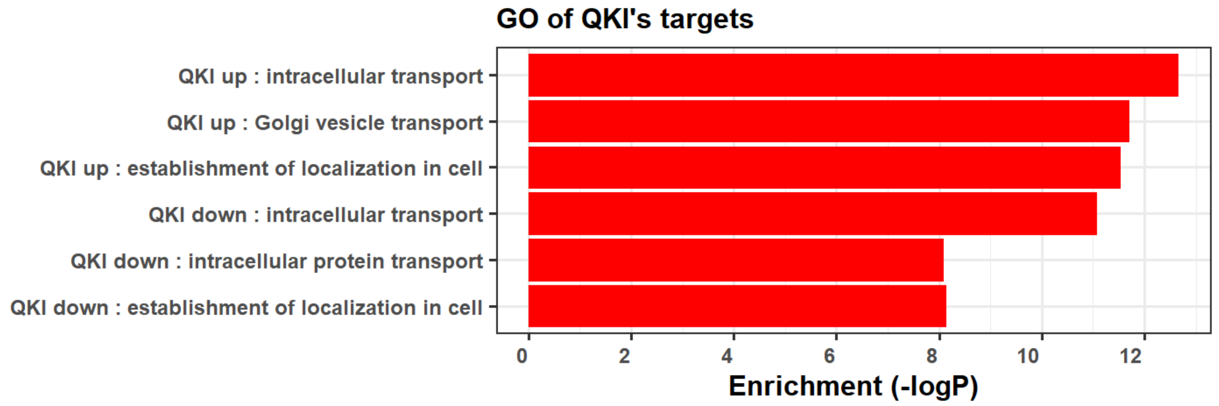
